# Temperature and dissolved organic matter shape marine prokaryotic activity and gene expression in a sub-arctic sea

**DOI:** 10.1101/2024.11.13.623445

**Authors:** Dennis Amnebrink, Ashish Verma, Daniel Lundin, Johan Wikner, Jarone Pinhassi

## Abstract

Temperature and dissolved organic matter (DOM) are important drivers of microbial activity, but their effects, alone or in combination, on the physiological responses of sub-arctic prokaryotic assemblages remain poorly understood. In a northern Baltic Sea one-month mesocosm experiment, we therefore exposed a coastal microbial community to temperature and nutrient regimes representative of winter and early summer (i.e., 1°C and 10°C, with and without DOM additions) in a 2×2 factorial design. Midway through the experiment, specific growth rates were highest for the 10°C plus DOM treatment (TN; ∼2.5 day^-1^), comparable for the 1°C plus DOM (N) and the 10°C (T) treatments at ∼1.0 day^-1^; and low for the control (1°C, no DOM enrichment [C]; 0.2 day^-1^). Taxonomic analysis of metatranscriptomes uncovered broad treatment specific responses, and a PERMANOVA on the 182,618 transcribed genes revealed statistically significant effects of both temperature and DOM, and significant interaction effects between the two (altogether involving 18% of genes). Significant differences in transcription identified by EdgeR analysis included *Nitrosopumilus* genes for ammonium uptake and ammonia oxidation in the 1°C mesocosms (C, N), membrane transporters for small organic acids in the N-treatment, genes for nitrogen and phosphorus assimilation along with molecular chaperones in the T-treatment, and dominance of Oceanospirillales genes for energy and growth metabolism in the TN-treatment. These metatranscriptomic responses were associated with changes in e.g. prokaryotic growth rates and growth efficiency, providing clues to how successional changes in community composition and metabolism are directed by temperature and DOM as central factors underlying environmental change.

**Importance:** It is recognized that increases in temperature and dissolved organic matter loading are key to understanding how climate change will influence polar ecosystems. Still, little is known of the effects of these factors on the physiological responses of Arctic prokaryotes. Since prokaryotes are principal drivers of biogeochemical cycles, we investigated how temperature and dissolved organic matter influence prokaryotic transcriptional responses in taxonomy and metabolic pathways. The metatranscriptomics analyses uncovered broad treatment specific responses linked with changes in prokaryotic community composition, growth rates, and growth efficiency. Yet, and importantly, the expression of most metabolic functions remained stable, suggesting a pronounced functional resilience of the prokaryotic community enabled by shifts in the dominance of different taxa. This emphasizes the large potential and importance of identifying the metabolic functions that underlie the divergence of prokaryotic communities in response to environmental changes projected to alter some of the most vulnerable marine environments.

## Introduction

Heterotrophic prokaryotes are key players in biogeochemical cycling, which can be attributed to their capability to metabolically transform dissolved organic matter (DOM) (1). In temperate marine ecosystems, the composition and function of prokaryotic communities exhibit pronounced seasonality (2, 3). In winter, coastal temperate and Arctic communities tend to exhibit elevated abundances of ammonia-oxidizing Archaea and/or Betaproteobacteria (4, 5). Next, following the onset of the productive season characterized by the spring phytoplankton bloom, bacteria that take advantage of bloom-derived organic matter flourish (6, 7). This can include members of the Flavobacteriia, opportunistic Gammaproteobacteria, and the Roseobacter clade (2, 8). In contrast, the summer season witnesses a surge in microbial community members adapted to low-nutrient conditions (9, 10). These changes in the premises for microbial growth and the associated development of microbial community composition imply that microbially mediated element fluxes will vary on seasonal scales.

Temperature, inorganic nutrients, and dissolved organic matter are important drivers structuring the composition and functionality of marine microbial communities (11–15). Moreover, bacterial growth and community composition dynamics depend to a large extent on the concentration and quality of DOM (14, 16–18). DOM concentrations and temperature typically covary in the Baltic Sea, reaching peak levels during the summer season when primary production peaks. Inorganic nutrient concentrations, in contrast, are high in winter and low in summer (19). The fact that environmental drivers covary, whether positively or negatively, makes teasing apart their individual impacts on microbial communities a complex endeavor. Yet, manipulations of environmental drivers in micro-or mesocosm scale experiments is an effective approach for elucidating distinct effects of such drivers on microbial communities (20, 21).

Metatranscriptomics is a powerful approach to determine the responses of marine microbial communities to environmental changes (22–25). More precisely, gene expression analyses can identify the molecular underpinnings of the strategies microbial communities employ to deal with alterations in growth conditions mediated by environmental drivers. This includes transcriptional changes in substrate acquisition genes and metabolic pathways to changing nutrient availability (23, 26, 27). Moreover, in some cases, metatranscriptomics has been successfully used to link the regulation of single genes to large scale metabolic fluxes of elements, such as the *amoA* gene in ammonia oxidation (28). In the context of ocean acidification, metatranscriptomics has uncovered shifts in the expression of genes related to pH homeostasis and stress in marine microorganisms (20). These findings demonstrate how metatranscriptomics can inform about reactions of marine microbes to environmental shifts and give valuable understanding of the mechanisms shaping community responses that determine carbon and nutrient cycling dynamics.

Verma et al. (2023) recently carried out a mesocosm experiment in the northern Baltic Sea with treatments including incubations at 1°C and 10°C, with and without DOM additions, in a 2×2 factorial design (29). The experiment started in early March to allow maintaining characteristics of the microbial community typical of winter (e.g. rates and composition) in this latitude and to induce characteristics mimicking those induced by early summer conditions in temperature and DOM supply. Notably, prokaryotic specific growth rates (µ) increased to similar values in the single-treatment mesocosms with increased temperature or DOM, but remained low in the control. Yet, the community composition in these treatments was distinct from each other and from the control. The combination of increased temperature and DOM enrichment resulted in the highest *µ* and the largest change in community composition (29). To further understand these dynamics in microbial characteristics, we here set out to investigate how the prokaryotic transcriptional responses in taxonomy and metabolic pathways were influenced by temperature and DOM manipulations simulating winter as compared to early summer conditions in sub-Arctic waters. We also tested for statistical interaction effects between temperature and DOM on the transcriptional level. We hypothesized that the relative prokaryotic expression patterns would diverge such that DOM enrichment would select for distinct taxa and functions in the mesocosms maintained at in situ temperature (1°C) as compared to the mesocosms with a temperature representative of early summer (10°C).

## Methods

### Experimental setup and sampling

On March 5th 2020, twelve indoor mesocosms were filled with 2 m^3^ each of water from the Bothnian Sea at Umeå Marine Sciences Centre, Sweden (63°34’ N, 19°50’ E), using the facility inlet-system. In these northerly waters, the transition from winter to early summer temperatures occurs within two months from early April onwards (Pinhassi 2000). In addition, we considered it a benefit to start the experiment with a winter microbial community since it is characterized by low prokaryotic abundances and relatively high diversity (2, 3, 30).

Triplicate mesocosms were assigned as; control (C; 1°C [in situ temperature], no DOM addition), DOM addition (N; 1 °C, addition of yeast extract), elevated temperature (T; 10 °C, no DOM addition), and finally elevated temperature plus DOM addition (TN; 10 °C, addition of yeast extract). The DOM enrichments were done twice a week throughout the experiment immediately after each sampling, by adding 477 mg yeast extract (Merck Millipore, granulated, cat. no. 1.03753.0500) dissolved in 100 ml Milli-Q water per mesocosm. This resulted in a total enrichment of 59 µM C, 16 µM N, and 1 µM P (based on measures in the added yeast extract solution) over the course of the experiment. The addition of yeast extract was done to simulate increased release of cell components that make up labile DOM during early summer due to protist and zooplankton excretions and sloppy feeding. Light was set to a 12:12 h diurnal cycle, with gradual light change using LED lamps (lightDNA R258 and software lightDNA8, Valoya, Finland). Light cycle conditions corresponded to end of April conditions, with an average of 122 μmol m^-2^ s^-1^ on the surface. Sampling for standard microbial ecology variables was carried out twice a week and sampling to collect DNA and RNA was carried out weekly.

### Water chemistry and microbial variables

Details on sampling of water chemistry and microbial variables are presented in Verma et al. (2023). Briefly, prokaryotic abundance was determined by epifluorescence microscopy and prokaryotic growth was determined based on thymidine incorporation rates of ^3^H-labeled thymidine. For dissolved organic carbon, samples were filtered through sterile 0.22 µm filters, acidified, and stored in the dark at 4°C. Chlorophyll *a* samples were filtered onto Whatman 25 mm diameter GF/F filters, and the filters were incubated in 96% ethanol in the dark for 12 hours. The filters were pelleted by centrifugation at 3500 rpm for 10 min. The chlorophyll *a* concentration in the supernatant was estimated through fluorometry at excitations and emissions wave lengths of 433 nm and 673 nm, respectively (Luminescence spectrometer LS-55, Perkin Elmer).

### RNA sampling, extraction and sequencing

For the collection of metatranscriptomic samples, filtration using peristaltic pumps was carried out in climate rooms set to the mesocosm temperatures (1°C or 10°C). Water was first prefiltered through 47 mm diameter 3.0 µm polycarbonate filters and biomass was collected on 0.22 µm Sterivex filters (filtration lasted a maximum of 30 min), to which 1.8 ml RNAlater was added. Samples were stored at –80°C until extraction. The day 10 and day 17 samples were selected for metatranscriptomic analysis based on the specific growth rates in the different treatments (Fig. 1e), to provide two consecutive data points where the specific growth rates were fairly similar in each of the treatments yet captured principal differences between treatments across the experiment.

**Figure 1.**
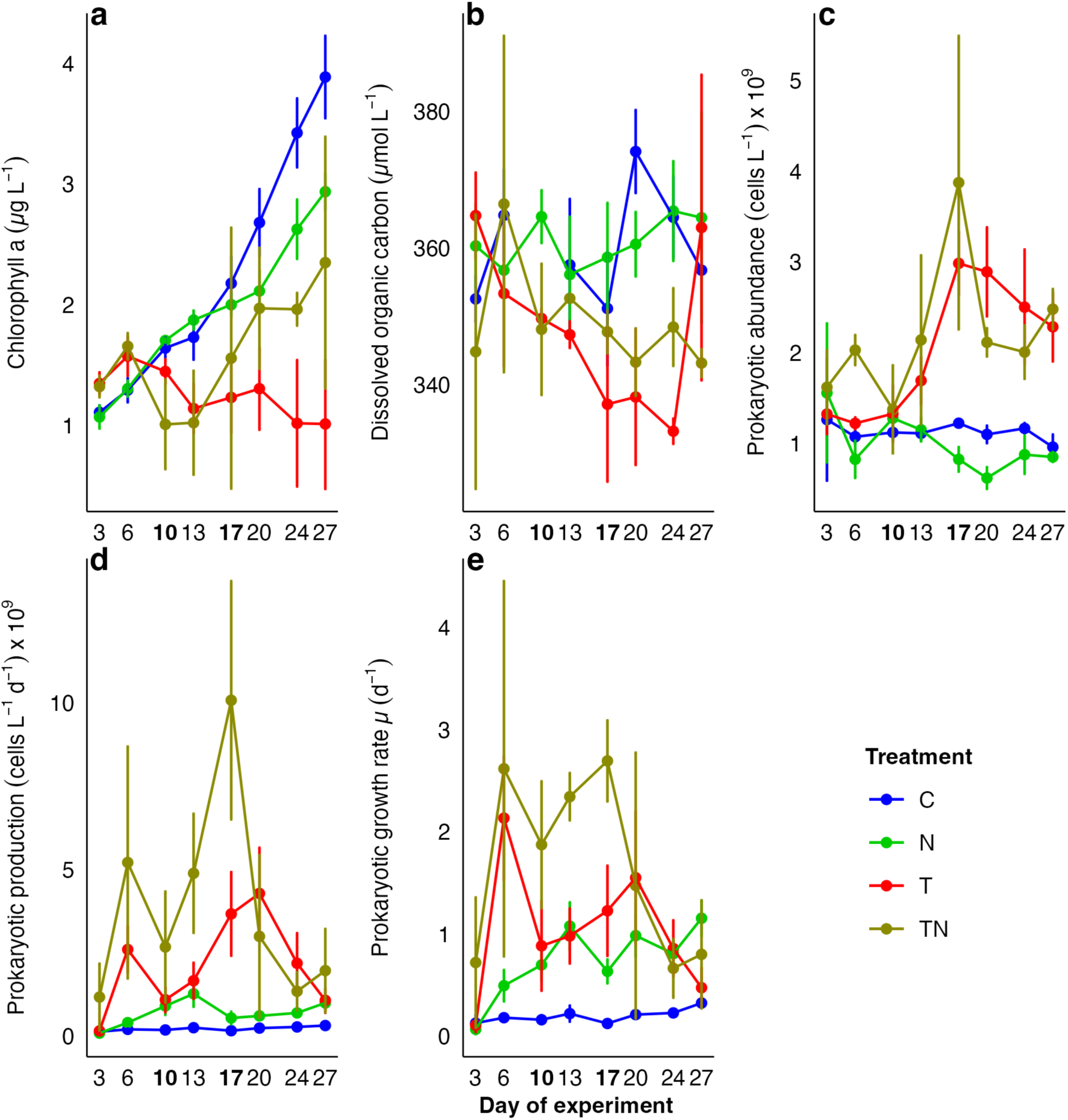
Temporal patterns of basic microbial ecology variables over the course of the one-month mesocosm experiment. Concentrations of (a) chlorophyll *a*, (b) dissolved organic carbon, (c) prokaryotic abundance, (d) prokaryotic growth, and (e) the specific prokaryotic growth rate. Error bars denote standard deviations (n=3; except for dissolved organic carbon where n=2 on day 10 in the C treatment). Days 10 and 17 in boldface indicate when analyses of metatranscriptomics were done. Panel c, d, and e redrawn from Verma et al. (2023).

Total RNA was extracted using an RNeasy Mini kit (Qiagen), following a protocol adapted from (31). Briefly, 0.22 µm Sterivex filters were thawed on ice and the RNAlater reagent was removed before proceeding to cut the filters with a razor. Filters were put in tubes with a 1 mL solution consisting of RLT-lysis buffer, β-Mercaptoethanol (10 µl/mL RLT buffer) and 2 spoons 200 µm Zirconium beads (OPS diagnostics). Cells were lysed using a vortex for 15 minutes, followed by centrifugation at 4°C for 5 min at 5000×g. The supernatants were transferred to tubes containing 1 volume 70% ethanol and mixed by repeated pipetting. Extraction was continued using the RNeasy mini kit by the manufacturer’s manual. Total RNA was eluted twice using 30 µL preheated RNase free water. Samples were DNAse treated using the TURBO DNA-free kit (ThermoFisher Scientific) following the manufacturer protocol. The RNA yield was measured using Qubit and Nanodrop. Before transcriptomic analysis, each sample was checked for genomic contamination using a 16S rRNA gene PCR and gel electrophoresis. rRNA was depleted using the RiboMinus transcriptome Isolation kit and RiboMinus Concentration Module (ThermoFisher Scientific). Ribosomal RNA molecules were then removed by binding to magnetic beads and only the rRNA-depleted fraction was further concentrated and purified using silica spin columns. RNA was amplified using the MessageAmp II-Bacteria RNA Amplification kit (ThermoFisher Scientific) following the manufacturer’s instructions. Lastly, final RNA was quantified and stored at –80**°**C until sequencing. Samples were sequenced on the NovaSeq6000 with 2×150 bp size at SciLifeLab (Stockholm, Sweden).

### Bioinformatic pipeline and data analysis

Samples were run with the nf-core framework (Ewels et al., 2020) using a prerelease version (“https://github.com/LNUc-EEMiS/metatdenovo” [shrivelled_shaw] DSL2 – revision: 83f8d3ec80 [dev]”) of the nf-core/metatdenovo assembly pipeline (https://nf-co.re/metatdenovo, Di Leo et al. in prep.). Briefly, samples were quality controlled with and trimmed using Trimgalore (Ver 0.6.6, https://zenodo.org/records/7598955), FastQC (Ver 0.11.9, (32)) and MultiQC (Ver 1.8, (33)). Adapter removal was carried out using cutadapt (Ver 3.2, (34)). Sequences were assembled using Megahit (Ver 1.2.9, (35)). Open reading frames (ORFs) were identified with Prodigal (Ver 2.6.3, (36)) and functionally annotated with eggNOG-mapper (evolutionary genealogy of genes: Non-supervised Orthologous Groups, version 2.0.8-2, (37)). For functional analyses, we chose to sum over the column named seed_eggNOG_ortholog in the eggNOG output. Taxonomy of the ORFs was assigned against phyloDB using EUKulele (Ver 1.0.4, (38)). The number of ORFs called, annotation success and number of reads mapping to ORFs, for each sample were then compiled from the output files (Table S1). Taxonomic and functional annotations of ORFs are in Table S2. The summary of larger pathways was done by grouping the KEGG-pathways based on the larger categories they constituted.

The resulting output was imported into R Ver 3.6.3 (39), where non-prokaryotic transcripts were filtered out before further analysis. A principal component analysis (PCA) on eggNOG annotated functions was carried out on Hellinger transformed raw counts, and a Permutational multivariate analysis of variance (PERMANOVA) with the eggNOGs was performed using the Vegan package (Ver 2.5-7, (40)). For comparison of gene expression categorized into larger metabolic pathways, open reading frame counts (ORFs) were normalized to transcripts per million (tpm) per sample to compensate for different library sizes and different ORF lengths, then averaged per treatment and time point. To analyze the effects of temperature, DOM and their interaction, a pairwise differential gene expression analysis was performed using statistical models with EdgeR (Ver 3.26.8, (41)) on raw counts of eggNOGs. To be included in the analysis, eggNOGs needed to pass a filtering step of a threshold of 15 counts in total and to be present in at least two samples. To correct for multiple testing, P-values were adjusted using the false discovery rate (fdr) correction. To be considered differentially abundant between treatments, a cutoff fdr-value of <0.05 was used. Compared treatment pairs with EdgeR were N-C, T-C, and TN-T to identify the effect of temperature and the effect of DOM addition at different temperatures. Lastly (TN-T)-(N-C) was used as the statistical model to determine the interaction effects of temperature and DOM.

## Results

### Basic microbial ecology variables

The Chl *a* concentration steadily increased during the 27 days of the experiment in the 1°C mesocosms, reaching 3.8 and 2.9 µg dm^-3^ in the C and N treatments, respectively. In contrast, in the 10°C mesocosms (T and TN), Chl *a* decreased until day 13, after which it steadily increased in TN but remained low in T throughout the experiment (Fig. 1a). Dissolved organic carbon (DOC) concentrations were fairly stable at ∼350 µM through the experiment in the 1°C mesocosms, but tended to decrease in the 10°C mesocosms (Fig. 1b). Pairwise analysis showed that the DOC concentrations were significantly different between the 1°C and 10°C mesocosms (Wilcoxon signed rank test, p<0.05, n = 24).

Prokaryotic abundance was highest in the 10°C mesocosms (T, TN) reaching peak abundance on day 17 (Fig. 1c), followed by a gradual and sharp decline in the T– and TN-treatment, respectively. Prokaryotic production followed a similar pattern, with sharp declines in the 10°C mesocosms from days 17 and 20 onwards (Fig. 1d). The specific growth rate (*µ*) increased strongly in the 10°C mesocosms from day 3 to 6 (Fig. 1e), then remained high around 2.2 d^-1^ in TN, while it dropped to ∼1.2 d^-1^ in T until day 17 and 20. In the N-treatment, *µ* gradually reached 1.0 d^-1^ on day 13, and remained at this level for the rest of the experiment. The C mesocosms showed low and stable *µ* (∼0.2 d^-1^). Prokaryotic abundance, production, and Chl *a* were found to be significantly different among the different pairwise treatments (Wilcoxon signed rank test, p<0.01, n = 24), except the T-TN comparison for prokaryotic abundance and Chl *a*.

### Overall metatranscriptomics analysis

Principal component analysis (PCA) of the prokaryotic community transcription on day 10 and 17 showed a clear separation between treatments (Fig. 2a). Treatments belonging to different temperature regimes separated on the primary axis, while the secondary axis separated the treatments with and without dissolved organic matter (DOM) additions. Moreover, some separation between the day 10 and 17 samples was observed within each treatment.

**Figure 2.**
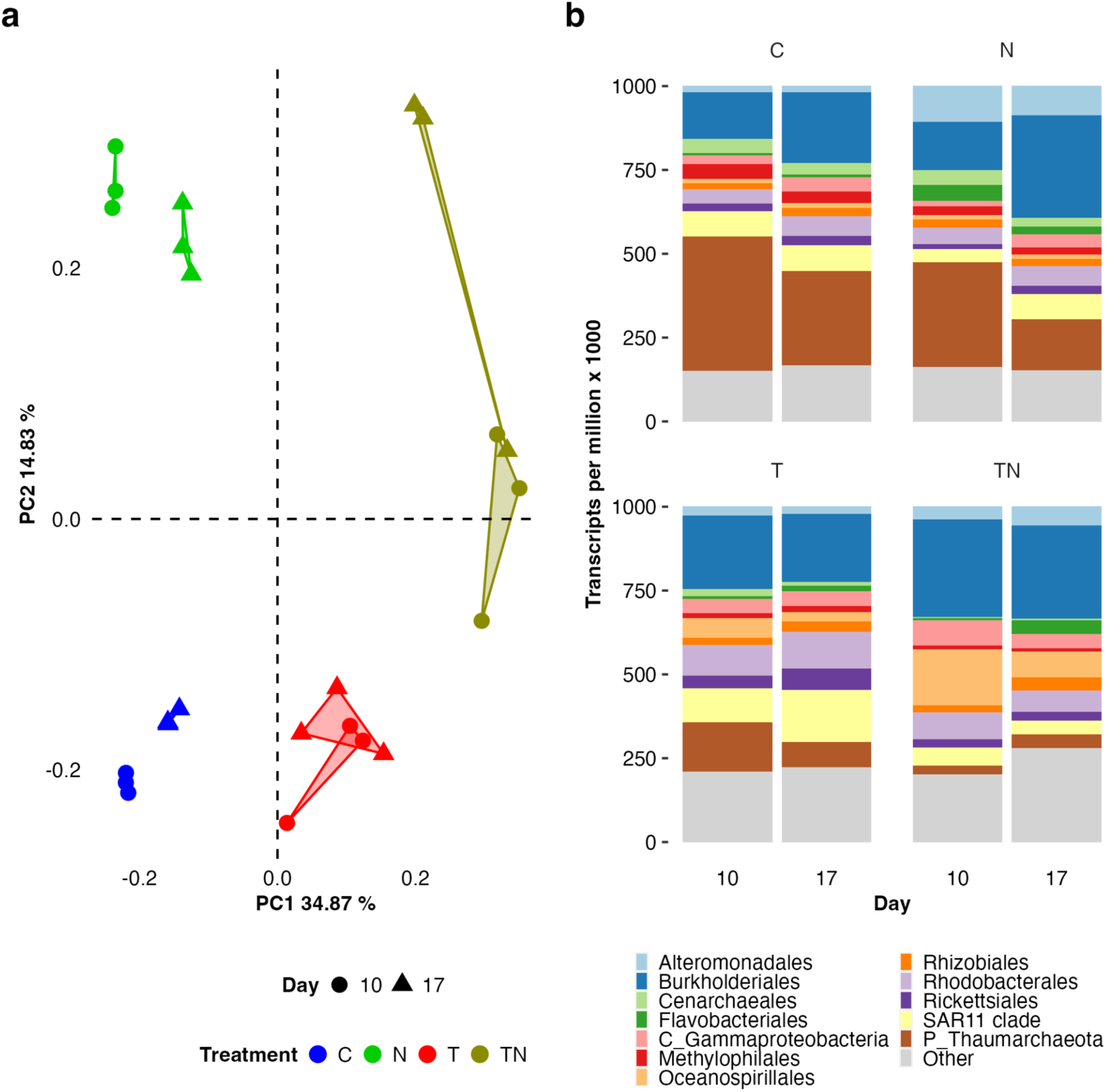
Comparison of relative prokaryotic expression in the mesocosms on days 10 and 17. (a) PCA-plot of relative expression based on the eggNOG-annotation system, and (b) order or higher-level taxonomic annotation of functional expression; in the legend, prefix C and P denote class or phylum respectively, no prefix denotes order-level.

Analysis of the taxonomic affiliation of the transcripts showed that the 1°C mesocosms (C, N) on day 10 were largely dominated by Archaea (up to 45%), which remained high also on day 17 (less so in the N-treatment; Fig. 2b). The grand majority of archaeal expression was accounted for by the genus *Nitrosopumilus* (Fig. 3a). Burkholderiales made up 10-25% of the transcription in all treatments (Fig. 2b), with a higher relative transcription of *Rhodoferax ferrireducens* in the 1°C mesocosms (C, N), while the 10°C mesocosms (T, TN) instead showed a higher fraction of the family *Oxalobacteraceae* (Fig. 3b). The DOM enriched treatments (N, TN) showed elevated expression of Flavobacteriales and Alteromonadales (Fig. 2b), with the most responsive Alteromonadales being *Shewanella frigidimarina* NCIMB 400 and *Shewanella denitrificans* OS217 (Fig. 3d). Expression in the T-treatment was characterized by alphaproteobacterial orders, where Rhodobacterales and SAR11 reached relative expression levels of 8-20% (Fig. 2b). The most notable feature of the expression in the TN-treatment was the development of the gammaproteobacterial order Oceanospirillales, which reached its highest relative expression on day 10 (17%; Fig. 2b). Oceanospirillales expression was accounted for by several genera and species in the T-treatment, and by *Marinobacterium stanieri* and *Neptuniibacter caesariensis* in the TN-treatment (Fig. 3c). Taxa with lower relative transcription levels, yet involved in ecologically potentially important processes, are presented in Table S2, such as *Nitrospira* spp. that typically contributes to nitrification.

**Figure 3.**
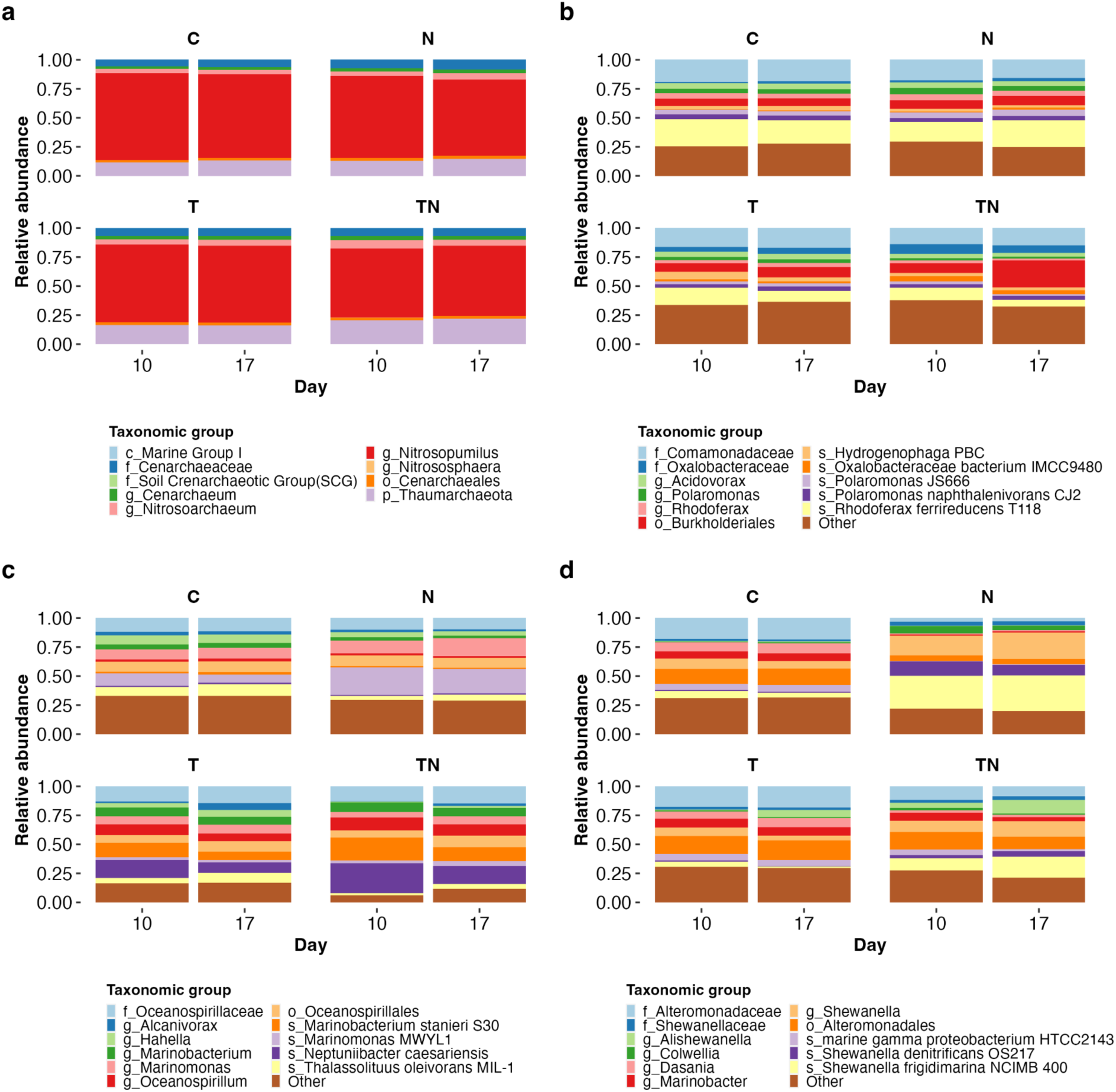
The distribution of relative expression within selected taxonomic groups. (a) the phylum Thaumarchaeota, (b) the order *Burkholderiales*, (c) the order *Oceanospirillales*, and (d) the order *Alteromonadales*. In the legends, prefixes denote taxonomic level; “s_” = species, “g_” = genus, “f_” = family, “o_” = order, “c_”= class, “p_” = phylum.

The most abundantly transcribed prokaryotic pathways in the KEGG database on both day 10 and day 17 were *Translation* and *Energy metabolism* (Fig. 4a). The *Translation* pathway relative expression levels were about two-fold higher in the 10°C than in the 1°C mesocosms on day 10 (which had evened out until day 17). The four KEGG pathways *Cell motility*, *Nucleotide metabolism*, *Signal transduction*, and *Membrane transport* reached higher relative expression in the 10°C mesocosms on day 10, and the latter two peaked in the T-treatment on day 17. Strikingly, the ammonium transport gene *amt*, represented by the manually added “Ammonia transport” category in Fig. 4a, showed roughly ten-fold higher relative expression in the C mesocosms compared to the other mesocosms on both day 10 and day 17. A tendency to higher levels of “Ammonia transport” also in the T mesocosms can be noted – resulting in the lowest relative expression levels being observed in the treatments with DOM additions (Fig. 4a).

**Figure 4.**
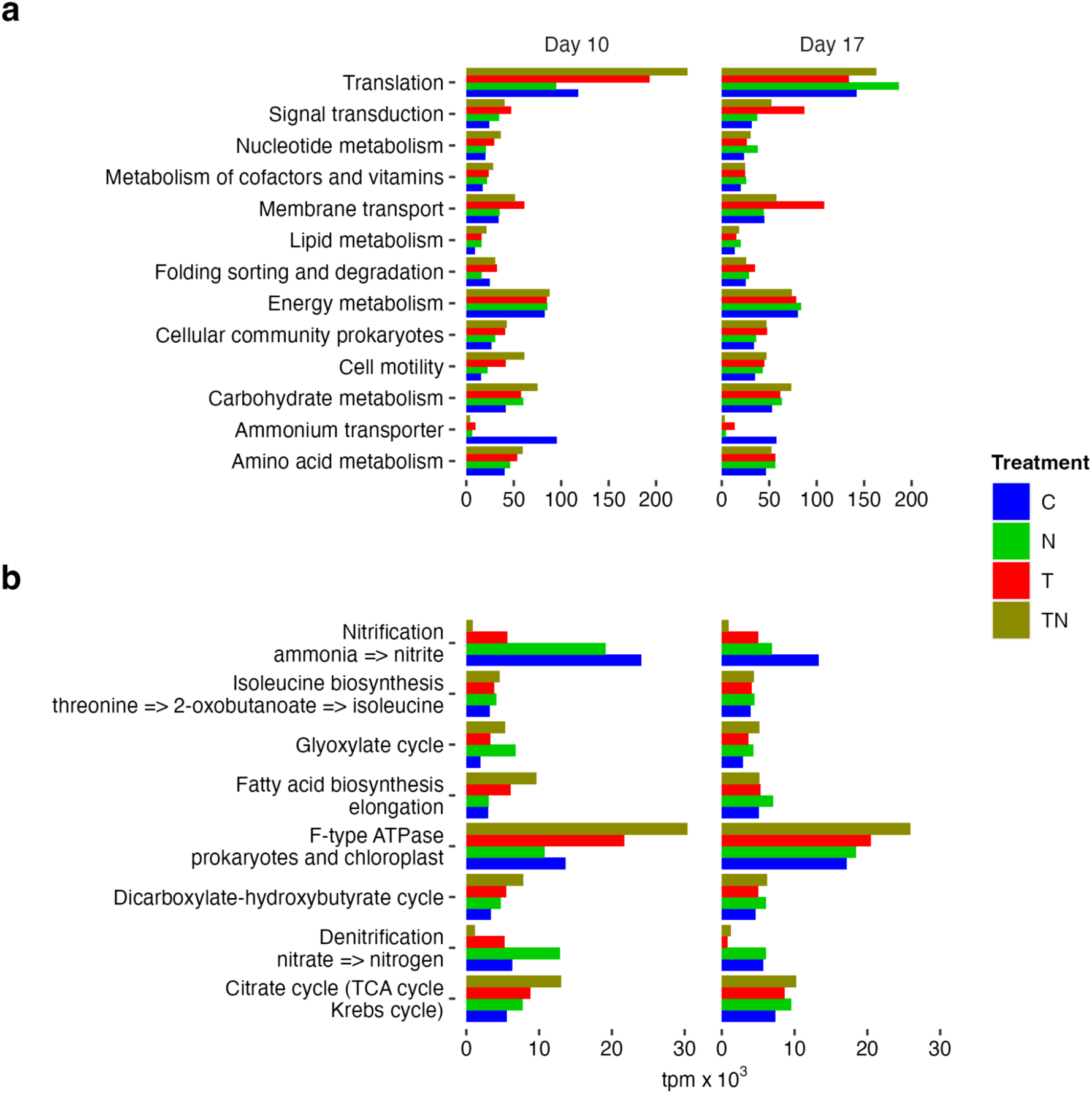
Relative expression values of dominant prokaryotic metabolisms in the mesocosms on days 10 and 17 according to the Kyoto Encyclopedia of Genes and Genomes (KEGG) database. (a) KEGG pathways, (b) KEGG modules. The “Ammonium transporter” category in panel a was added manually and is not part of the KEGG metabolic pathways. Note differing x-axis scales in the upper panels (a) compared to the lower panels (b).

At the specific module level of the KEGG classification (Fig. 4b), the most abundant module was *F-type ATPase*, which showed higher relative expression in the 10°C mesocosms (especially in TN on day 10). At somewhat lower expression levels, the *Fatty acid biosynthesis* and *Citrate cycle* modules had similar expression patterns as the *F-type ATPase module* on day 10. The *Nitrification* module reached high relative expression in the 1°C mesocosms (C and N), especially on day 10, whereas *Denitrification* in these mesocosms were five-fold higher than in the 10°C mesocosms on day 17. The *Glyoxylate cycle* module tended to reach higher relative abundance in the DOM-amended treatments (N and TN) (Fig. 4b).

### Statistically differentially abundant transcripts

A PERMANOVA based on the 2×2 factorial design of the mesocosm experiment and the 182,618 transcribed eggNOGs on day 10 revealed significant effects of both temperature and DOM, in agreement with the clustering of samples in the PCA (p<0.001, Fig. 2a). Moreover, the PERMANOVA showed a significant interaction effect between temperature and DOM (p<0.005, Table S3). The subsequent edgeR analysis to identify the extent of these transcriptional responses showed that up to ∼10% of the eggNOGs in any of the tested contrasts were significantly differentially transcribed (p<0.05; Table S4). Accordingly, and focusing first on the responses of DOM addition at the two temperatures, the statistical analysis of transcription in the mesocosms maintained at 1°C (i.e. the N versus C contrast) showed that 424 and as many as 18,202 eggNOGs reached significantly higher relative expression in the C– and N-treatment, respectively. The corresponding analysis for the 10°C mesocosms (i.e. the TN versus T contrast) showed that 3,868 and 2,156 eggNOGs reached higher transcription in the T– and TN-treatment, respectively. Instead focusing on the effect of increased temperature alone (i.e. the T versus C contrast), 6,227 eggNOGs were found to be transcribed at significantly higher relative abundance in the 10°C (T) treatment. The interaction effect of temperature and DOM resulted in 3,776 eggNOGs where the factors had a negative interaction (i.e. significantly influenced by the interaction in C) and 330 eggNOGs in the positive interaction (i.e. significantly influenced by the interaction in TN).

### Taxonomy and function of differential transcripts

We next classified the significantly differentially expressed eggNOGs into major metabolic KEGG categories (i.e. pathways) subdivided by taxonomy (Fig. 5). This showed first and foremost a pronounced dominance of “Ammonium transporter” expression in the 1°C controls, accounted for by the archaeal genus *Nitrosopumilus*. In the other mesocosms, a majority of the transcription of the significantly expressed eggNOGs was associated with the four KEGG pathways *Carbohydrate*, *Energy*, and *Amino Acid metabolism* plus *Translation*. However, there were pronounced differences in the taxonomy of the prokaryotes to which these metabolisms were associated in the different treatments. Accordingly, in the N-treatment, there was a pronounced overrepresentation in relative expression of Alteromonadales among the significant eggNOGs as compared to the total expression (Fig. 5). Also Flavobacteriales were represented in a broad variety of KEGG pathways in the N-treatment, whereas *Nitrosopumilus* were abundant only in the *Energy Metabolism* pathway (encoding functions involved in carbon fixation and ammonia oxidation among others) (Table S4). In the T-treatment, there was a particularly high expression of *Translation* as compared to the other metabolic categories, mediated mainly by Burkholderiales, Oceanospirillales and Rhodobacterales (Fig. 5). The SAR11 clade accounted for more of the differentially expressed eggNOGs in the T-treatment than in any of the other treatments. A key feature of the differentially expressed eggNOGs in the TN-treatment was the dominance of Oceanospirillales in the majority of the KEGG pathways.

**Figure 5.**
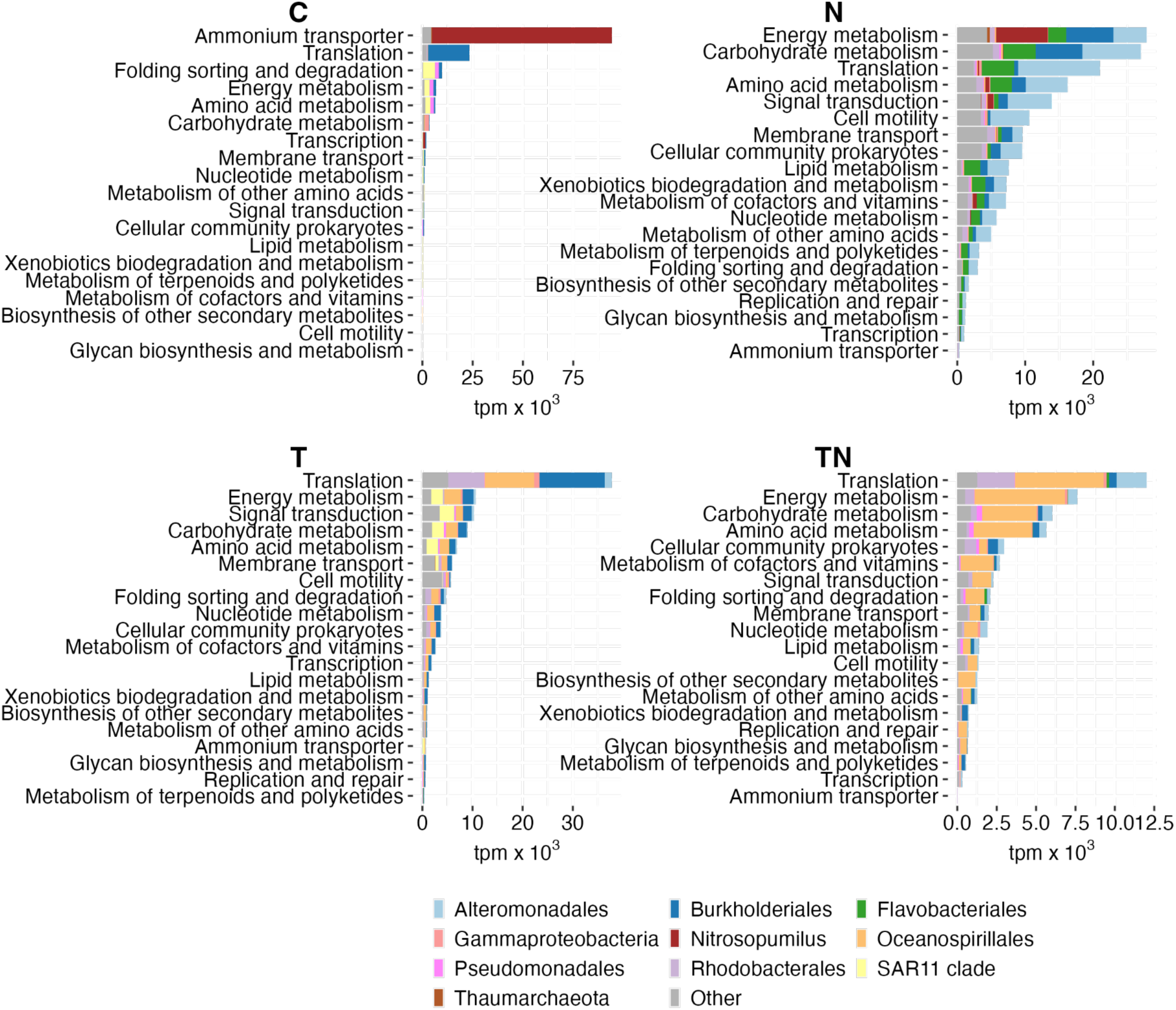
Distribution of significantly differentially expressed genes (eggNOGs) in KEGG categories and by order level taxonomy on day 10. The 1°C control mesocosms (C), the 1°C plus DOM treatment (N), the 10°C treatment (T), and the 10°C plus DOM treatment (TN). The “Ammonium transporter” category in panel a was added manually and is not part of the KEGG metabolic pathways.

To gain more specific insight into metabolisms influenced by temperature and DOM, we visualized the 30 significant eggNOGs with highest relative abundance (Fig. 6). This showed that in the C mesocosms, the vastly dominant gene was *amtB* encoding the *Nitrosopumilus* ammonium transporter. Also notable were the SAR11 clade orthologs of *groL*, which encode a key stress response chaperone protein that assists in the turnover of misfolded proteins, and the peroxidoredoxin gene, which is involved in protection against oxidative stress. A majority of the remaining eggNOGs encoded subunits of Burkholderiales ribosomes.

**Figure 6.**
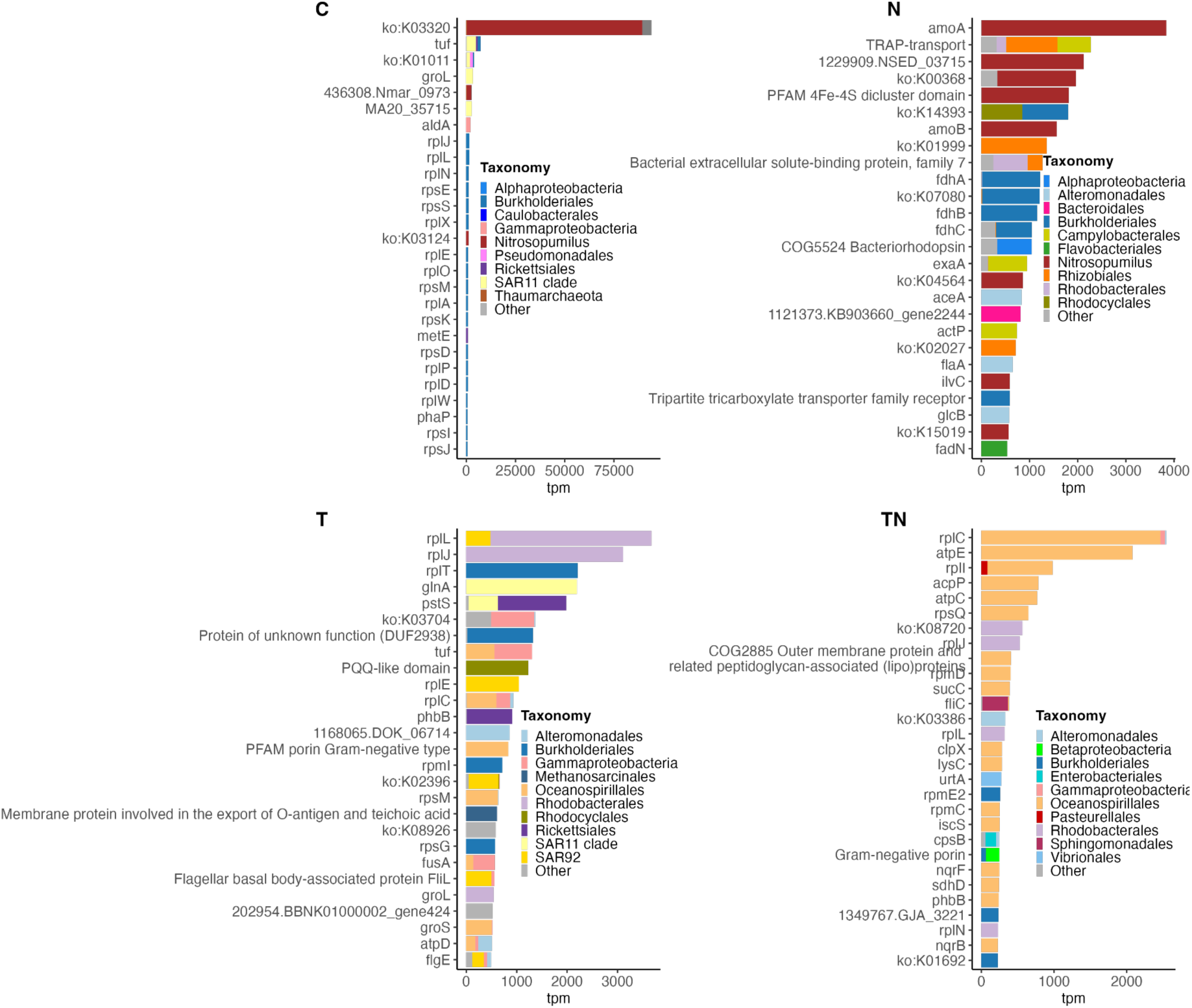
Visualization of the 30 most abundant significantly differentially expressed genes (eggNOGs) in the mesocosms. The 1°C control mesocosms (C), the 1°C plus DOM treatment (N), the 10°C treatment (T), and the 10°C plus DOM treatment (TN).

In the N treatment, five different *Nitrosopumilus* eggNOGs were among the most highly expressed (Fig. 6), most notably *amoAB* encoding the two archaeal subunits for ammonia monooxygenase that allows obtaining energy from ammonia oxidation. These were accompanied by *nirK*, encoding the nitrite reductase enzyme that forms nitric oxide (NO) as intermediate in either of the two proposed pathways of nitrite formation (42) – the end product of *Nitrosopumilus* ammonia oxidation. Regarding substrate utilization of Bacteria, several taxa showed high relative expression for TRAP membrane transporters involved in organic acid utilization. Also alphaproteobacterial proteorhodopsins (PRs) allowing photoheterotrophy were highly expressed in the N treatment. Curiously, the eggNOGs for the two key anaplerotic enzymes in the glyoxylate shunt of the TCA cycle, isocitrate lyase (AceA) and malate synthase (GlcB), were both highly expressed (Fig. 6) – which allow growth on e.g. C2 compounds like pyruvate.

The most highly differentially expressed eggNOGs in the T-treatment were two Rhodobacterales ribosomal proteins (Fig. 6), consistent with this taxon reaching their highest overall relative expression in this treatment (Fig. 2b). The high relative abundance of both *glnA* and *pstS* showed that especially the SAR11 clade bacteria made substantial transcriptional investments in both nitrogen and phosphorus assimilation (Fig. 6). Glutamine synthetase (GlnA) plays a key role in nitrogen assimilation by catalyzing the reaction glutamate plus ammonia to generate glutamine, which acts as nitrogen donor in several metabolic pathways. PstS is the high-affinity substrate-binding subunit of the phosphate ABC-type membrane importer. Whereas proteorhodopsin genes were found in the N-treatment, photoheterotrophy in the T-treatment was represented by the *pufA* gene, encoding the alpha-subunit of light-harvesting complex I (involved in pigment-binding) in bacteria carrying out aerobic anoxygenic photosynthesis with bacteriochlorophyll *a*. The chaperone genes *groL* and *groS* were both among the top 30 genes in the T-treatment. Lastly, in the TN-treatment, Oceanospirillales genes dominated among the top 30 genes with particularly high relative expression of genes for ribosomal proteins (prefix *rp*) and ATP synthase subunits (*atpCE*) (Fig. 6).

### Statistically significant interaction effects between temperature and DOM

Statistical analysis showed that the relative expression levels of 2.2% (n=4,106 genes) of the total of 182,618 transcribed genes were significantly influenced by interaction effects between temperature and DOM (edgeR; p<0.05). The analysis identified 3,776 genes for which the additive effects of the two factors resulted in a lower relative transcript abundance compared to either factor alone, here termed as negative interaction. Conversely, the transcription of 330 genes showed additive effects between the two factors that resulted in a higher relative transcript abundance compared to either factor alone, here termed as positive interaction. Fig. 7a shows the relative expression levels and taxonomic assignment of the genes with negative (left panel) and positive interactions (right panel). Note the dominance of *Nitrosopumilus* genes under the negative interaction (Fig. 7a, left panel), and the reciprocal distribution patterns for the different Proteobacteria under different interactions (left versus right panel). Unfortunately, the majority of *Nitrosopumilus* eggNOGs did not obtain a functional classification category, and were thus left as not assigned (NAs) (Fig. 7b).

**Figure 7.**
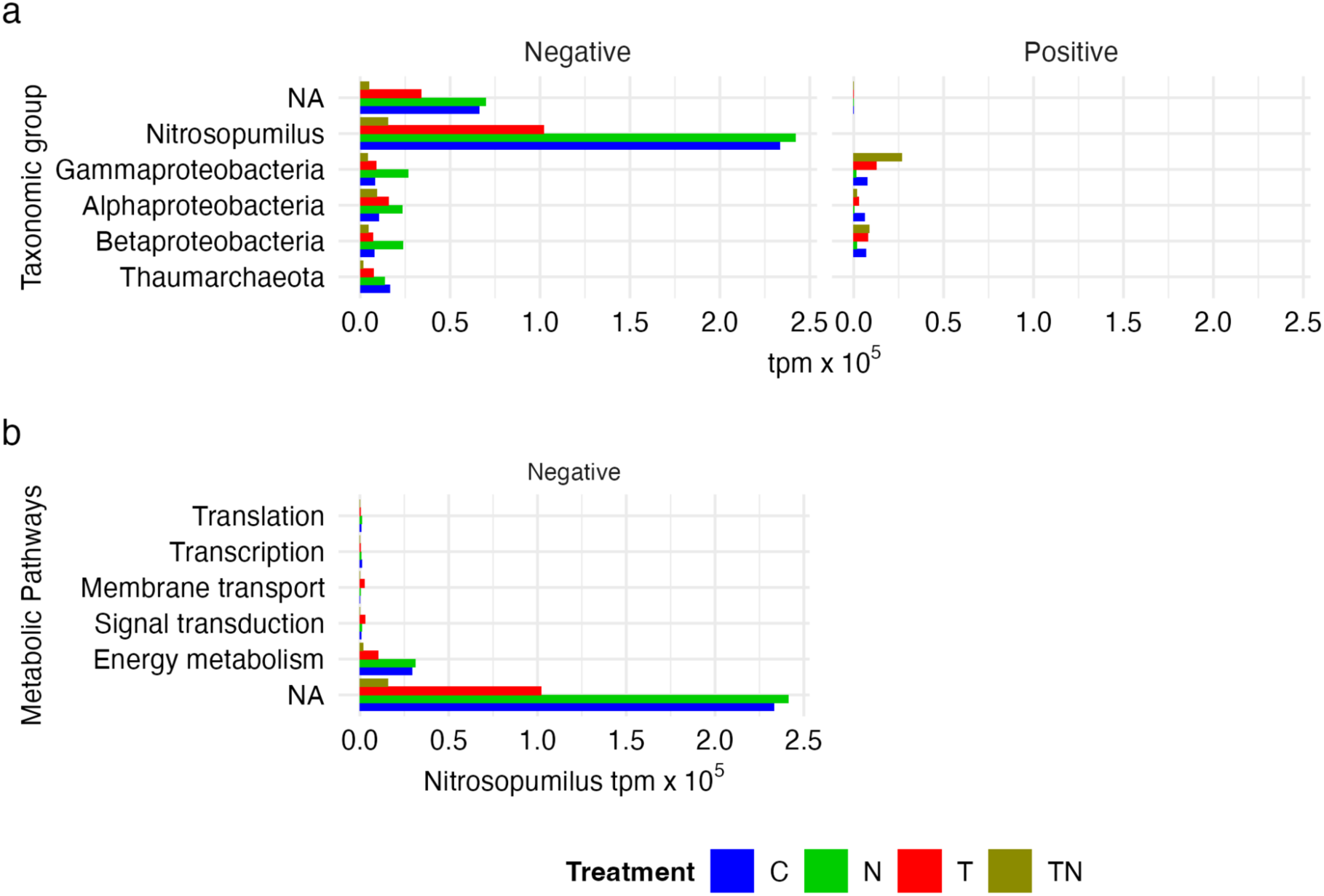
Distribution of transcripts for genes with a significant interaction effect by DOM addition and temperature manipulation on day 10. (a) Relative transcription levels for the taxa with highest relative transcription; “Negative” denotes eggNOGs under influence of negative interaction effects between temperature and DOM, “Positive” denotes eggNOGs with positive interaction. (b) Relative transcription levels of the most prominent *Nitrosopumilus* KEGG pathways (subset from panel 7a “Negative”).

## Discussion

The current work builds on an experiment aimed at investigating how prokaryotic community composition and activity are influenced by manipulation of temperature and DOM to simulate the transition from winter to early summer characteristics in a subarctic coastal sea (29). Our current metatranscriptomics analysis of functional genes in samples collected midway through the experiment (days 10 and 17) aligned with the findings based on 16S rRNA gene amplicon analysis, with samples clustering primarily by treatment and next by time. The divergence in community composition and transcriptional patterns in the N– and T-treatments were particularly interesting, in light of the similarity in community growth rates (∼1.0 day^-1^) in these two treatments on day 10 and 17. This implies that transcriptional adjustments together with community turnover are important to uphold critical prokaryotic community characteristics (at times referred to as ecosystem functions; here exemplified by specific growth rate) in the face of diverging environmental conditions.

Although temperature and DOM availability are each long-recognized drivers of microbial activities and succession, few studies have made direct comparisons of the relative importance of the two. Divergence in metabolic strategies as a function of temperature has for example been observed in an analysis of a major set of cultured prokaryotic isolates in renowned culture collections, showing that growth temperature selects for a variety of adaptations in enzyme functions (43). Regarding DOM, the chemical characteristics and availability of different compounds is recognized to have a strong influence on both the activity and transcription of different prokaryotic taxa (14, 44–46). This supports that our observed shifts in community composition were associated with shifts in metabolism or possibly, reciprocally, that the metabolic characteristics of different taxa contributed to induce changes in community composition.

The forcing on the prokaryotic community that was induced by manipulation of temperature and DOM was considerable and was chosen to represent highly distinct growth conditions – winter and early summer in a subarctic coastal sea. Indeed, pronounced changes were observed both in the community composition (29) and in the taxonomic affiliation of the expressed functional genes as seen here. Nevertheless, over the analyzed time course (days 10 and 17 for metatranscriptomics), several taxa remained abundant, and their transcriptional activity remained similar, albeit at altered relative levels. This agrees with the general observation that the manipulations had statistically significant effects on only 18% of the functional genes (out of the total 182,618 genes; here using “genes” to refer to the eggNOGs). These observations can be compared with the findings of Liu et al. (2022) that a core metabolic network is shared between highly remote microbiomes (Mount Everest and the Mariana Trench) even though the microbiomes are very different taxonomically (47). We think it is important to keep in mind the authors’ concluding wording that “primary metabolic functions could be always conserved”. In our reading, this provides inspiration for considering the taxonomic affiliation of metabolic functions to aid ecological interpretations of distribution patterns of genes and transcripts in relation to environmental drivers.

Before discussing treatment-specific prokaryotic responses in the mesocosm experiment, we wish to highlight the broad distribution and abundance of Burkholderiales (Proteobacteria). We find noteworthy the high and fairly even relative expression levels (up to ∼25%) of Burkholderiales in all mesocosms (and up to over 50% of prokaryotic 16S rRNA gene amplicons; (29)). Especially given the pronounced divergence in overall transcription patterns and the accompanied shifts in dominant prokaryotic taxa between treatments. Some of this variation could be partially explained by differential responses of distinct taxa within the order between treatments, such as *Oxalobacteraceae* spp. in the 10°C mesocosms or *Oxalobacteraceae* bacterium IMCC9480 in the TN-treatment. Despite their high relative expression levels, Burkholderiales accounted for only minor portions of the statistically verified differences in expression, particularly in the TN-treatment. These findings suggest that Burkholderiales were successful through a generalist life history strategy, as previously proposed (22). It is important to point out here that, in general, order level taxa often include multiple families and many genera, so that as in the case of Burkholderiales they may range from generalist opportunists to specialized oligotrophs (as shown for different populations by (17); for detail on oligotrophic Burkholderiales, see (48)). Burkholderiales are known to be abundant in winter waters in the Baltic Sea (2, 49) and accounted for up to 75% of the prokaryotic community in a Baltic Sea subarctic estuary during spring (17). Yet, their ecological roles in the Baltic Sea remain largely unexplored.

The elevated relative expression of Archaea – especially *Nitrosopumilus* – in the 1°C mesocosms (C, N) is in line with the observation of Archaea in 16S rRNA gene amplicon data during winter in the Baltic Sea (2). At our chosen levels of manipulation, the archaeal transcriptional signal appeared to be more sensitive to elevated temperature than DOM enrichment, implying low temperature as an important factor for Archaea to be competitive in Baltic Sea surface waters. Curiously, these findings are in line with previous studies emphasizing the distribution of elevated nitrogen and carbon metabolisms of ammonia oxidizing *Nitrosopumilus* in substantially warmer surface waters and coastal habitats (50, 51). This includes the NW Mediterranean Sea (50), where it should be noted that the minimum temperatures in winter are usually typically ten degrees higher than the winter temperature we encountered (i.e. similar to the “elevated” temperature we used). The most prominent feature of *Nitrosopumilus* metabolism is its ammonia oxidizing capability, from ammonia to nitrite – whereby the (little) energy and reducing power gained are used for fixation of CO_2_ through a 3-hydroxypropionate/4-hydroxybutryrate pathway variant (rendering them autotrophic) (52, 53). Indeed, higher transcripts of *amoA* have been correlated with higher potential nitrification rates in winter waters (4). Experimental work with *Nitrosopomilus maritimus* strain SCM1 shows that it carries out relatively high ammonia oxidation rates even under extremely oligotrophic conditions, thanks to a high specific affinity for reduced nitrogen (54).

Alongside *Nitrosopumilus* transcription, but at much lower relative levels, we found *Nitrospira* sp. transcription that indicated comammox – the complete oxidation of ammonia to nitrate (55, 56) potentially occurred in our experiment. The *Nitrospira* genus has been suggested to be adapted to oligotrophic conditions (57, 58). Similar to the *Nitrosopumilus* distribution in our experiment, the higher relative expression of *Nitrospira* sp. in the 1°C mesocosms (C, N), and stronger decline in the N-treatment as compared to the C-treatment, further strengthens the notion that they are adapted to nutrient limiting conditions. Previously, higher abundances of comammox bacteria have been shown in a coastal environment (59), suggesting that *Nitrospira* sp. together with *Nitrosopumilus* contribute to nitrogen transformation during winter conditions in the Baltic Sea.

The prominent statistically supported transcriptional response in the N-treatment of the Alteromonadales and Flavobacteriales showed that these were highly competitive taxa in winter waters (1°C) when provided with labile but primarily complex DOM. It should be noted though that these taxa reached an elevated proportion of the total community transcription also with DOM at 10°C (TN-treatment). Alteromonadales occurs during phytoplankton spring blooms in the North Sea where their glycosyl hydrolases could contribute to the utilization of polysaccharides (8, 60). Similarly, a metaproteomics analysis of microbial plankton in a NW Atlantic bay (Bedford Basin, Halifax) showed that Alteromonadales reached their highest proportion of the community (4%) in the winter to spring transition (61). Moreover, laboratory experiments with natural prokaryotic communities have uncovered that enrichment with high-molecular-weight DOM (45) and polysaccharides or nucleic acids (14) triggers increased proportions of Alteromonadales expression. Also Flavobacteria are recognized as potent utilizers of polymeric substrates (as those found in yeast extract) with enzymes to assimilate and metabolize proteins and polysaccharides (62), such as glycosyl hydrolases and peptidases (63, 64). Additional significant genes of potential ecological importance were the several TRAP transporters transcribed by different taxa that indicated small organic acids were an important resource. Interestingly, while proton-pumping rhodopsins are considered to primarily be of use to photoheterotrophic bacteria under carbon-limited conditions (65), proteorhodopsin genes were significantly expressed in the DOM-enriched treatment (N). In this context it is notable that the TCA cycle glyoxylate shunt gene transcription could potentially be linked with both the TRAP transporter and the rhodopsin expression, since the glyoxylate shunt has been tightly linked to the use of small organic acids and rhodopsin light harvesting (66), although a direct coupling between these genes was not observed in their taxonomic affiliation in the current study.

The increase in temperature to 10°C from the *in situ* temperature of 1°C had a similar effect on prokaryotic growth rates as the DOM enrichment. Yet, instead of triggering the transcription by taxa recognized for their opportunistic lifestyle, temperature led to higher relative expression levels of Rhodobacterales and SAR11 clade bacteria, often associated with phytoplankton and oligotrophic conditions, respectively. The Rhodobacterales includes the *Roseobacter* clade that can constitute up to 20% of cells in coastal environments (67) and shows a large diversity in their genomic content – ranging from streamlined genomes adapted to oligotrophic conditions to larger genomes encoding a large metabolic flexibility (68, 69). Despite this diversity, elevated temperature (likely in combination with lower resource availability) rather than DOM seemed to be the main driving factor of Rhodobacterales expression in our study. In particular, this raises the question if or how higher temperatures may influence the available substrate pool by lowering the activation energy for catabolic pathways involved in the utilization of natural DOM pools. Likewise, the elevated relative expression of SAR11 in the T-treatment was in line with their adaptation to oligotrophic growth conditions (70). But it is also in agreement with the finding that higher temperature favors the growth of slow-growing prokaryotes, as was recently shown for data sets spanning seasons, latitude, and depth from different seas (10). In Abreu et al. (2023), the principal seasonal data were from the Linnaeus Microbial Observatory (LMO) in the Baltic Proper, where increasing temperatures in summer favored the growth of both SAR11 and other slow-growing taxa. The significant relative expression of the chaperone encoded by *groL* in the C and T mesocosms, could indicate that even low-nutrient adapted SAR11 clade bacteria and roseobacters are forced to struggle to maintain their cell integrity when DOM availability is low in these waters.

The highest community growth rates and growth efficiencies were observed in the TN-treatment, where the gammaproteobacterial order Oceanospirillales reached their highest relative abundance in transcription. The incidence of genes associated with growth among the genes with highest relative expression in the TN-treatment, such as ATP synthase and ribosomal proteins, suggested that this taxon had the capacity to opportunistically exploit the increased temperature, especially in combination with increased DOM availability. Positive responses of a variety of Gammaproteobacteria to elevated temperature have been observed in both field studies and laboratory experiments in the Baltic Sea ((21); and references therein) and mesophilic Oceanospirillales of the genus *Marinobacterium* have been isolated on nutrient replete medium and yeast extract (71).

The interaction effect of temperature and substrate concentrations on heterotrophic bacterial activity has been extensively studied, as reviewed by (72). Yet, results have been contrasting, and sometimes no interaction effect is evident. In our study, no interaction effect on prokaryotic rate measurements were recorded (29). However, the gene expression data complemented the traditional rate measurements such that several transcriptionally active taxa and their genetic functions showed marked changes due to the interaction effect. This was particularly the case for nitrogen related metabolic pathways, potentially influencing elemental nitrogen and carbon fluxes through *Nitrosopumilus* metabolism. Incidentally, expression rates of the ammonia monooxygenase gene have been related to nitrogen flux rates in marine environments (28). This highlights the importance of considering how factors that structure prokaryotic community composition and activity, such as temperature and DOM, interact to influence biogeochemical cycling properties.

### Conclusions

Taken together, our metatranscriptomics results show that temperature and DOM both have pronounced effects on the transcriptional activity of prokaryotic communities and that they can cause interaction effects not easily detected with traditional bulk community measurements used in aquatic microbial ecology. Furthermore, although the community composition and the dominance in relative transcriptional investment by different prokaryotic community components changed substantially, the expression of the majority of transcribed functions were maintained. This suggests a pronounced functional resilience of the prokaryotic community that was made possible through shifts in the dominance of different taxa. We interpret this as if the transcriptional activation of a fairly limited set of genes in particular pathways (i.e., up to 18% of the total of 182,618 eggNOGs) in taxa responsive to the studied environmental changes enabled the prokaryotic community to respond and adapt functionally to the experimentally induced shifts in temperature and DOM. Importantly, this was associated with changes in ecologically important characteristics of the bacterial communities, such as growth rates and growth efficiency. Collectively, our observations pinpoint some potential key features of the prokaryotic communities associated with winter and early summer conditions, and how successional changes are induced by environmental conditions linked with seasonality.

## Code and data availability statement

Code used to produce figures and tables in this article can be found at: https://github.com/dennisamne/PROMAC_metaT Sequences and are accessible at the European Nucleotide Archive (ENA) under project: PRJEB63211.

## Acknowledgements

We thank Camilla Karlsson and Sabina Arnautovic for their hard work in the laboratory. We acknowledge the help from the personnel at Umeå Marine Science Centre (UMF) with aiding in carrying out the mesocosm experiment. This work was supported by the Kempe Foundation grant SMK-185 (JW), the Swedish strategic research area programme EcoChange (JW, JP), and the Swedish research council VR (2015-04254; JP). The computations and data handling were enabled by resources in projects NAISS 2023/6-240 and NAISS 2023/22-601 provided by the National Academic Infrastructure for Supercomputing in Sweden (NAISS), partially funded by the Swedish Research Council through grant agreement no. 2022-06725.

## SUPPLEMENTARY TABLES

**Table S1**. Sequencing and annotation statistics for each sample from the mesocosm experiment. *Table S1 is available on figshare:* https://figshare.com/s/216538b6ce177dfd1705

**Table S2**. Taxonomic and functional annotations of ORFs. t3 and t5 in the timepoint column denote samples from Day 10 and Day 17, respectively. *Table S2 is available on figshare:* https://figshare.com/s/7cfef141d37ae72c45f4

**Table S3**. Permutational multivariate analysis of variance (PERMANOVA) on the effects of temperature and DOM and their interaction on the gene expression (eggNOGs). *Table S3 is available on figshare:* https://figshare.com/s/8867bc11b40d9efa0b7d

**Table S4**. EdgeR statistics, identity, and taxonomic annotation of genes (eggNOGs) identified as significantly differentially transcribed. The contrast column indicates which treatments that were tested against each other. *Table S4 is available on figshare:* https://figshare.com/s/cd0d2e0ff96822f29add

